# Epithelial cell biomarkers are predictive of response to biologic agents in Crohn’s disease

**DOI:** 10.1101/2020.05.20.106518

**Authors:** Mark T. Osterman, Kelli L. VanDussen, Ilyssa O. Gordon, Elisabeth M. Davis, Katherine Li, Kate Simpson, Matthew Ciorba, Sarah C. Glover, Bincy Abraham, Xueyan Guo, Eric U. Yee, Felicia D. Allard, Jacqueline G. Perrigoue, Brian Claggett, Bo Shen, Thaddeus S. Stappenbeck, Julia J. Liu

## Abstract

**Objective:** Therapeutic efficacy of biologics has remained at about 50% for 2 decades. In Crohn’s disease (CD) patients, we examined the predictive value of an epithelial cell biomarker, ileal microvillar length (MVL), for clinical response to ustekinumab (UST) and vedolizumab (VDZ), and its relationship to another biomarker, intestinal epithelial cell (IEC) pyroptosis with respect to response to VDZ.

**Design:** Ileal biopsies from the UNITI-2 randomized controlled trial were analyzed for MVL as a predictor of clinical response to UST. In a 5-center academic retrospective cohort of CD patients, ileal MVL was analyzed to determine its predictive value for response to VDZ. Correlation between ileal MVL and IEC pyroptosis was determined, and the discriminant ability of the combination of two biomarkers to VDZ was examined.

**Results:** Clinical response in UST was significantly higher than placebo (65% vs. 39%, p=0.03), with patients with normal MVL (>1.7 µm) having the greatest therapeutic effect: 85% vs. 20% (p=0.02). For VDZ, clinical response with MVL of 1.35-1.55 µm was 82% vs. 44% (<1.35 µm) and 40% (>1.55 µm) (p=0.038). There was no correlation between ileal MVL and IEC pyroptosis. The combination criteria of ileal pyroptosis < 14 positive cells/1000 IECs or MVL of 1.35-1.55 µm could identify 84% of responders and 67% of non-responders (p=0.001).

**Conclusions:** Ileal MVL was predictive of response to UST and VDZ in prospective and retrospective CD cohorts. It was independent of ileal IEC pyroptosis, combination of the two biomarkers enhanced the discriminate ability of responders from non-responders to VDZ.

## Introduction

Biologics are antibodies targeting specific inflammatory pathways that have been used for the treatment of moderate-to-severe inflammatory bowel disease (IBD), including Crohn’s disease (CD) and ulcerative colitis (UC), for over two decades.^1-5^ There are several classes in clinical use, including anti-tumor necrosis factor (TNF),^1-3^ anti-integrin,^4, 5^ and anti-interleukin 12/23 (IL-12/23) antibodies.^6, 7^ IBD is a heterogeneous group of diseases with highly variable disease phenotypes and clinical courses, and patients will likely have different responses to each biologic class depending on their specific disease subtype. Despite widespread clinical adoption of anti-TNF therapy, nearly half of the CD patients placed on biologics do not experience clinical response, resulting in dramatic increase in healthcare costs without significant improvement in outcomes ^8, 9^ The annual costs of IBD treatment have increased by 30% within the last 5 years,^9^ and IBD-related hospitalizations quadrupled from 1998 to 2015.^10, 11^

Over the past decade, advances in disease pathogenesis from genomics, proteomics, and microbial and host metabolomic studies, also known as “multi-omics” approaches have identified key factors responsible for ongoing mucosal inflammation in IBD.^12-14^ The majority of effort to date has focused on the discovery and validation of biomarkers that can predict response to anti-TNF therapy. Mucosal and blood biomarkers that can predict responses to anti-TNFs using the multi-omics approach include oncostatin M,^15^ TNFR2 and IL13RA2, ^16, 17^ and TREM1.^18, 19^ However, integrating them into routine clinical care has not yet occurred.

We have recently discovered microvillar gene expression to play a role in the pathogenesis of mucosal inflammation and altered in CD patients.^20^ Moreover, a previously identified mucosal biomarker, called ileal microvillar length (MVL), was found to be a good cellular readout of the gene expression data, and ileal MVL was shown to be reduced in patients with active CD but increased with ustekimumab therapy.^20^ However, its predictive value for response to biologic therapies is unknown.

We also recently demonstrated that a novel mucosal biomarker, ileal intestinal epithelial cell (IEC) pyroptosis, an inflammatory form of cell death in the intestine, was predictive of clinical response to vedolizumab therapy in CD patients.^20^ In that multicenter retrospective cohort study, 89% of vedolizumab-treated patients with an ileal IEC pyroptosis level < 14 positive cells per 1,000 IECs achieved clinical response.

Given the above, the current study aimed to ascertain the predictive value of ileal MVL for therapeutic response to two biologics agents, ustekinumab and vedolizumab, in CD patients. Additionally, the correlation between the 2 epithelial cell biomarkers, ileal MVL and ileal IEC pyroptosis, in vedolizumab-treated CD patients was calculated. Finally, the utility of combining these biomarkers in order to discriminate responders from non-responders to vedolizumab in patients with CD was evaluated.

## Methods

### Study design and population

This study utilized 2 different cohorts, both prospective and retrospective, to address the various aims. To assess the predictive value of ileal MVL for response to ustekinumab in CD, a prospective cohort was used. This cohort consisted of patients enrolled from American study sites of the pivotal UNITI-2 randomized controlled trial of ustekinumab for CD.^6, 7^ For the purpose of this study, patients with active ileal or ileocolonic disease who had ileal biopsies obtained were included.

For all other aims which involved vedolizumab-treated CD patients, specifically investigating the association between ileal MVL and response to vedolizumab, correlating ileal MVL and ileal IEC pyroptosis, and determining the utility of combining these biomarkers to differentiate responders from non-responders to vedolizumab, a multicenter retrospective cohort was used. This cohort consisted of CD patients from 5 IBD centers across the United States (Cleveland Clinic Foundation, Houston Methodist Hospital, University of Florida, University of Pennsylvania, and Washington University). Patients aged 18-80 years with a confirmed diagnosis of CD based on standard clinical, radiological, endoscopic, and histological criteria who had been prescribed vedolizumab for moderate-to-severe CD, disease flare, corticosteroid dependence, failure of other biologic therapies, or allergies or adverse reactions to other agents, were included. Patients were required to have undergone colonoscopy with ileal biopsies prior to the initiation of vedolizumab. Individual patient charts were reviewed in the electronic medical records to obtain all clinical information. The study protocol was reviewed and approved by the Institutional Review Board at each study site.

### Outcomes

For the aim of evaluating the predictive value of ileal MVL for response to ustekinumab using the prospective randomized cohort, the following outcomes were assessed at 8 weeks post-induction: clinical response, defined as a reduction of Crohn’s Disease Activity Index (CDAI) of ≥ 100 points from pre-treatment baseline; clinical remission, defined as CDAI < 150; and endoscopic response, defined as a ≥ 50% reduction from baseline Simple Endoscopic Score for Crohn’s Disease (SES-CD) score. The primary outcome was clinical response to ustekinumab stratified by pre-treatment ileal MVL, using a threshold of 1.7 µm, which corresponds to 75% of the mean ileal MVL of the healthy general population.^20^ Pre-specified secondary outcomes included correlations between pre-treatment ileal MVL and baseline clinical activity, as measured by the CDAI, and baseline endoscopic activity, as measured by the SES-CD.

For all other aims involving vedolizumab-treated CD patients from the multicenter retrospective cohort, the following outcomes were evaluated approximately 6 months after initiation of therapy: clinical response, defined as a reduction of Harvey-Bradshaw Index (HBI) of ≥ 5 points from pre-treatment baseline; and clinical remission, defined as HBI < 5. The primary outcome was clinical response to vedolizumab by pre-treatment ileal MVL. The correlations between pre-treatment ileal MVL and baseline HBI and baseline SES-CD were also calculated.

### Assessment of ileal MVL

One set of biopsies was prepared for H&E-stained tissue sections. Brightfield images of H&E-stained resection margins were acquired with an Olympus BX51 microscope equipped with UPlanFL 10X/0.30, 20X/0.50, 40X/0.75, and 100X/1.30 Oil Iris objective lenses, an Olympus DP70 camera and DP Controller software or an Olympus DP22 camera and CellSens Standard v1.17 software. To quantify MVL and cell height, 5 enterocytes were measured per villus on 10 villi per sample, for a total of 50 cells measured per sample. Measured cells were visualized with the 100X objective lens and were located in the top one-third of a villus, where enterocytes have a matured brush border, in regions with well-oriented epithelial cells. We have previously demonstrated high inter-observer agreement with this method. ^20^

### Assessment of ileal IEC pyroptosis

Ileal biopsies collected from study patients during colonoscopy before the initiation of vedolizumab therapy were sectioned and stained for activated caspase stains using Maximus Biological Assay staining kits (Maximus Diagnostics LLC, Little Rock, AR) and anti-CD3 antibody to differentiate intraepithelial lymphocytes from IECs. Samples with at least 10 intact crypts/villi were analyzed for each patient to quantitate IEC pyroptosis by two blinded gastrointestinal pathologists (EUY and FDA). IECs were manually counted in 10 villi for the derivation of pyroptosis normalized per 1,000 IECs. Confocal images of the slides were acquired with Zeiss LSM 880 confocal microscope equipped with Airyscan (Zeiss USA, Dublin, CA).

### Sample size calculation

The sample size calculation was based on the assumption that there would be a difference in clinical response rate to ustekinumab of at least 30% between patients on therapy vs. placebo by ileal MVL cutoff value. A total of 84 patients (42 per group) would be needed to achieve 80% statistical power with a type I error (α) of 0.05, assuming equal numbers of patients in each group. Similarly, for vedolizumab response, we assumed that an ileal MVL cutoff or range could be identified that would provide a 30% difference in clinical response rate between patients at either ends of this cutoff or range. A total of 84 patients (42 per group) would be needed to achieve 80% statistical power with a type I error (α) of 0.05, assuming equal numbers of patients in each group.

### Statistical analysis

Baseline demographic, disease-related, and medication variables were collected and included, respectively: age, sex, race/ethnicity, and body mass index; disease phenotype, location, and activity (both clinical and endoscopic); and concomitant use of immunomodulators and corticosteroids, as well as history of prior anti-TNF exposure. Continuous variables were described using means with standard deviations or medians with interquartile ranges; categorical variables were expressed as numbers and proportions.

The probabilities of clinical response to ustekinumab and vedolizumab by pre-treatment ileal MVL were plotted graphically. Clinical and endoscopic outcomes stratified by ileal MVL thresholds or ranges were compared using Fisher exact tests. Likelihood ratio tests were used to assess for effect modification by ileal MVL. Spearman correlation coefficients were calculated to determine the relationships between ileal MVL or ileal IEC pyroptosis and baseline clinical and endoscopic activity. For prediction of clinical response to vedolizumab, an ileal IEC pyroptosis threshold of 14 positive cells / 1,000 IECs was used.^21^ The value of ileal MVL alone or in combination with ileal IEC pyroptosis for prediction of clinical response to vedolizumab was compared using Fisher’s exact test. All analyses were performed using STATA statistical software (StataCorp LP, College Station, Texas). Two-sided p-values < 0.05 were considered statistically significant.

## Results

### Response to ustekinumab in the randomized prospective cohort

For determination of ustekinumab response, a total of 106 CD patients from UNITI-2 (placebo = 36, ustekinumab = 70) biopsies were stained and analyzed, of which 95 patients (90%) had sufficient samples for analysis of ileal MVL. Baseline patient characteristics between placebo- and ustekinumab-treated patients were comparable (**Table 1**). The rate of clinical response was significantly higher in the ustekinumab-treated group: 65% (40/62) vs. 39% (13/33) (p = 0.03), with a similar trend in clinical remission rate of 42% (26/62) vs. 24% (8/33, p=0.12).

**Table 1:**
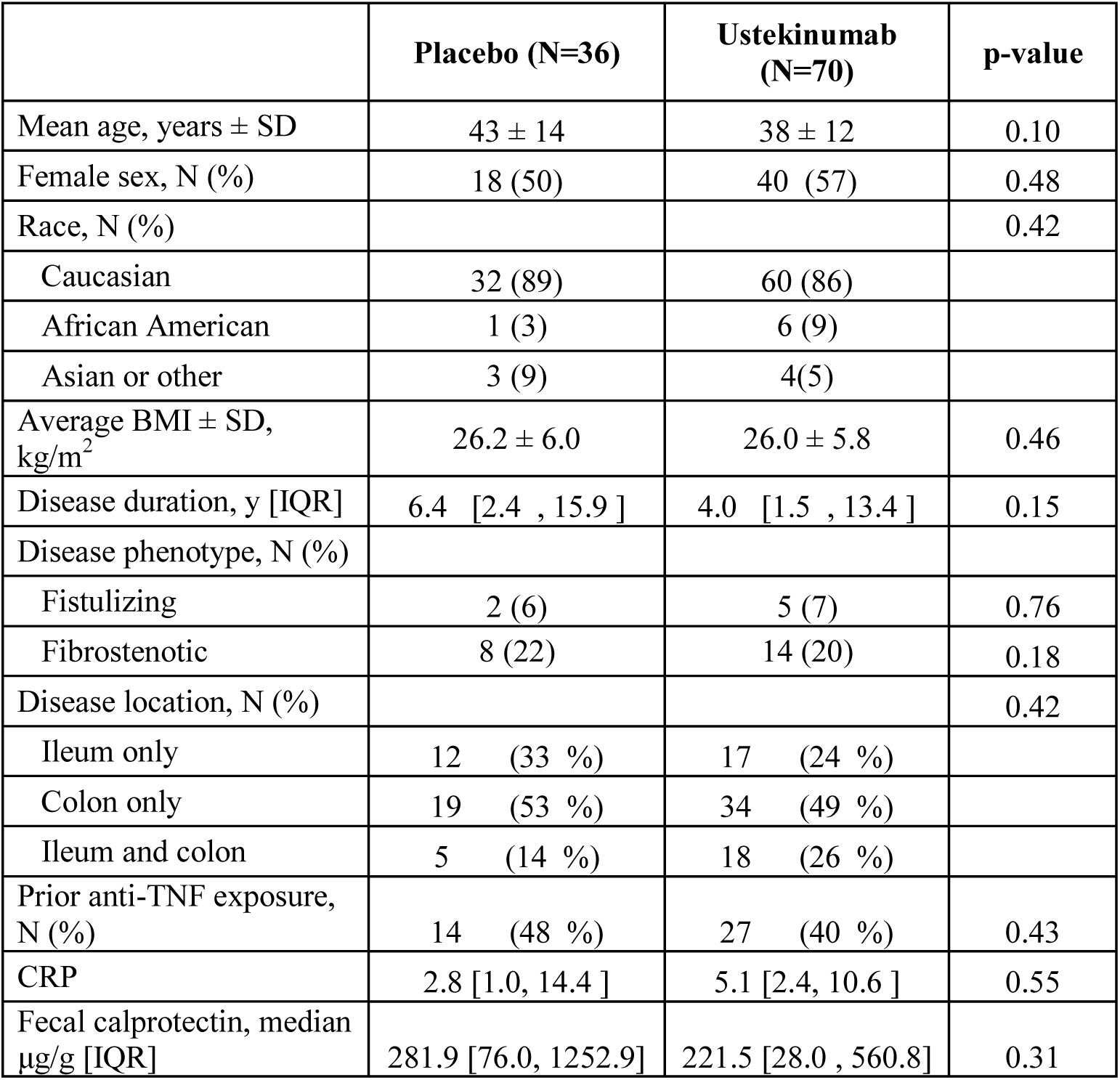
Baseline characteristics of the ustekinumab response cohort from UNITI-2.

|  | **Placebo (N=36)** | **Ustekinumab (N=70)** | **p-value** |
| --- | --- | --- | --- |
| Mean age, years ± SD | 43 ± 14 | 38 ± 12 | 0.10 |
| Female sex, N (%) | 18 (50) | 40 (57) | 0.48 |
| Race, N (%) |  |  | 0.42 |
| Caucasian | 32 (89) | 60 (86) |  |
| African American | 1 (3) | 6 (9) |  |
| Asian or other | 3 (9) | 4(5) |  |
| Average BMI ± SD, kg/m<sup>2</sup> | 26.2 ± 6.0 | 26.0 ± 5.8 | 0.46 |
| Disease duration, y [IQR] | 6.4 [2.4 , 15.9 ] | 4.0 [1.5 , 13.4 ] | 0.15 |
| Disease phenotype, N (%) |  |  |  |
| Fistulizing | 2 (6) | 5 (7) | 0.76 |
| Fibrostenotic | 8 (22) | 14 (20) | 0.18 |
| Disease location, N (%) |  |  | 0.42 |
| Ileum only | 12 (33 %) | 17 (24 %) |  |
| Colon only | 19 (53 %) | 34 (49 %) |  |
| Ileum and colon | 5 (14 %) | 18 (26 %) |  |
| Prior anti-TNF exposure, N (%) | 14 (48 %) | 27 (40 %) | 0.43 |
| CRP | 2.8 [1.0, 14.4 ] | 5.1 [2.4, 10.6 ] | 0.55 |
| Fecal calprotectin, median μg/g [IQR] | 281.9 [76.0, 1252.9] | 221.5 [28.0 , 560.8] | 0.31 |

As a continuous variable, ileal MVL was found to be an effect modifier of response to ustekinumab (p = 0.043), with the probability of response to ustekinumab having a positive linear relationship to ileal MVL with higher response rates as MVL increased, while placebo response had a negative linear relationship to ileal MVL with lower response rates as increased MVL (**Figure 1A**). Using the 75th percentile of ileal MVL in the healthy general population of 1.7 µm as a threshold, the greatest therapeutic effect and differences were seen in the normal MVL group (≥ 1.7 µm): clinical response rate of 85% (11/13) with ustekinumab vs. 20% (1/5) with placebo (Δ vs. placebo = 65%, p = 0.02) (**Figure 1B**); clinical remission rate of 62% (8/13) with ustekinumab vs. 0% (0/5) with placebo (Δ vs. placebo = 60%, p = 0.036) (**Figure 1C**). In the low MVL group (< 1.7 µm), rates were 59% (28/49) with ustekinumab vs. 46% (12/28) with placebo (Δ vs. placebo = 13%, p = 0.17) for clinical response (**Figure 1B**), and 37% (18/49) vs. 29% (8/28, Δ vs. placebo = 8%, p = 0.62) for clinical remission (**Figure 1C**). The therapeutic effect by endoscopic response, available in 71 patients, was also greater in patients with normal ileal MVL: 75% (6/8) vs. 20% (1/5), Δ vs. placebo = 55% (p = 0.053); compared to low MVL patients, with rates of 65% (24/35) vs. 48% (11/23), Δ vs. placebo = 17% (p = 0.11) (**Figure 1D**).

**Figure 1.**
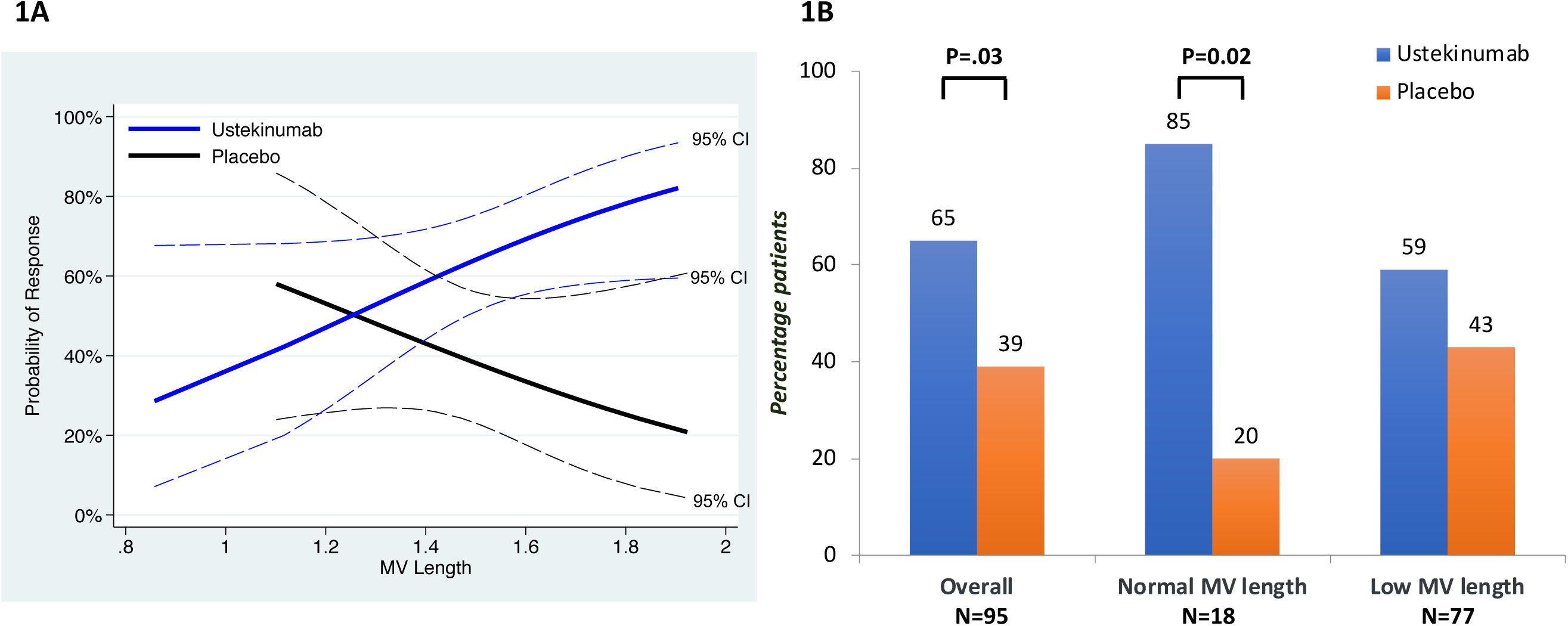
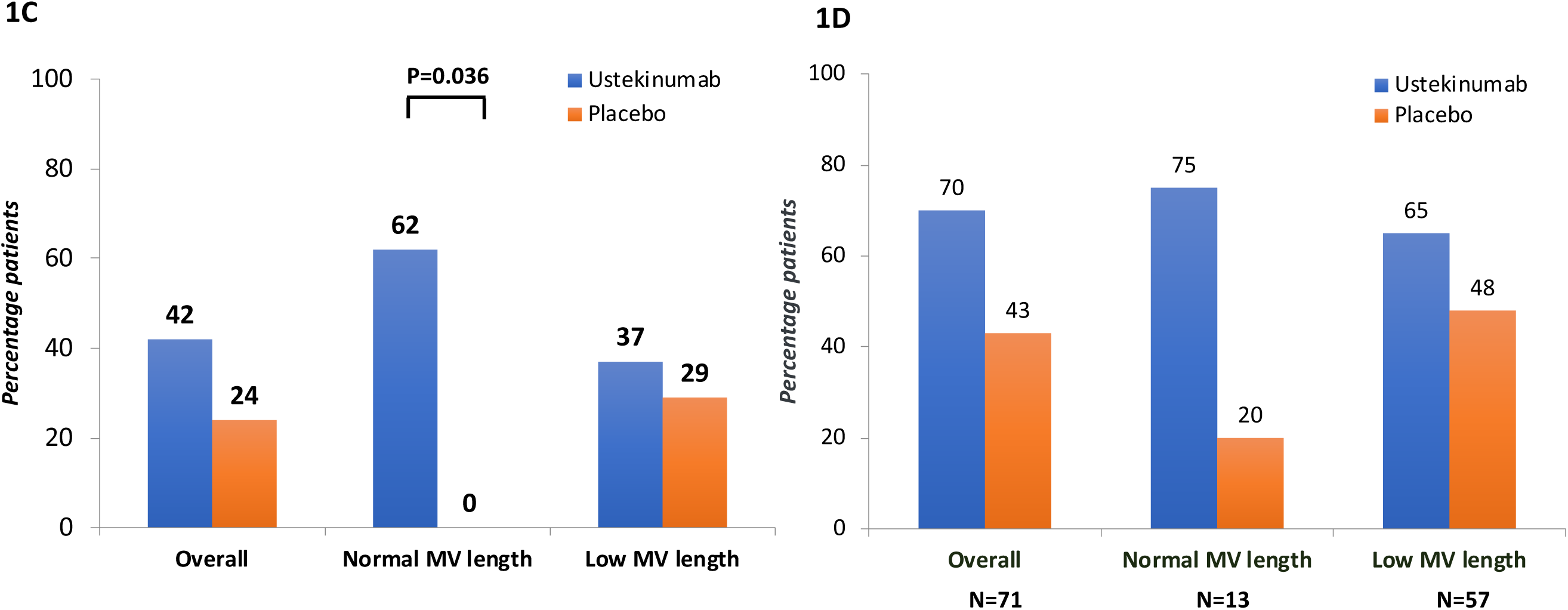
Ileal MVL is predictive of therapeutic response to ustekinumab in CD patients from UNITI-2. A) Ileal MVL is an effect modifier of clinical response to ustekinumab (N=95). B) Clinical response to ustekinumab stratified by pre-treatment ileal MVL (N=95). C) Clinical remission to ustekinumab stratified by pre-treatment ileal MVL (N=95). D) Endoscopic response to ustekinumab stratified by pre-treatment ileal MVL (N=71).

With respect to relationship with baseline disease activity, pre-treatment ileal MVL was not correlated with baseline clinical disease activity, as measured by CDAI (rho = −0.22) (**Figure 2A**). Ileal MVL was not correlated with baseline endoscopic disease activity by SES-CD (rho = −0.09) (**Figure 2B**).

**Figure 2.**
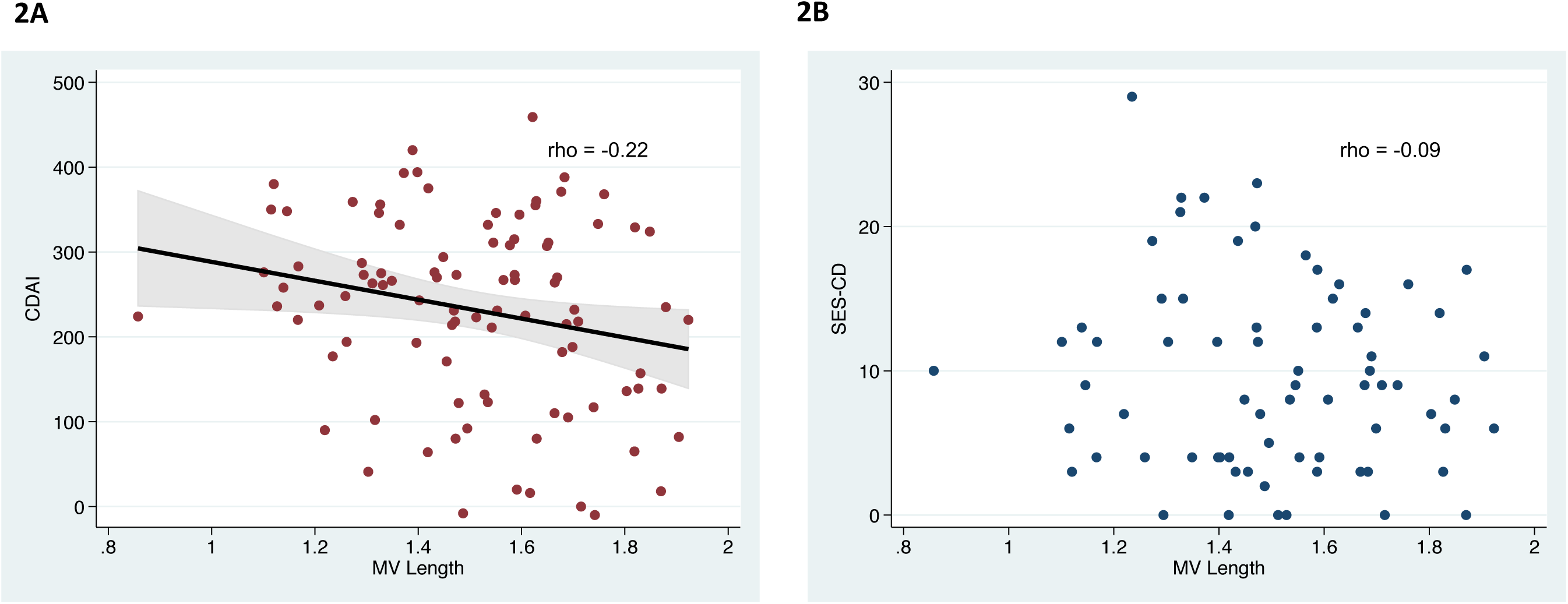
Correlation between ileal MVL length and baseline disease activity indices. A) Clinical disease activity by CDAI (N = 95). B) Endoscopic disease activity by SES-CD (N = 95).

### Response to vedolizumab in the multicenter retrospective cohort

For prediction of response to vedolizumab, 86 CD patients from the 5 IBD centers had ileal biopsies stained and analyzed, of which 64 had adequate samples for analysis of both ileal MVL and ileal IEC pyroptosis. The overall clinical response rate to vedolizumab was 59% (38/64). There were no significant differences in baseline patient characteristics, disease characteristics, concomitant medication use, or prior anti-TNF exposures histories between responders (N = 38) and non-responders (N = 26) (**Table 2**).

**Table 2:** Baseline characteristics of the vedolizumab response cohort from multiple academic sites.

|  | **Responders**<br>**(N=38)** | **Non-responders**<br>**(N=26)** | **p-value** | **Total**<br>**(N=64)** |
| --- | --- | --- | --- | --- |
| Mean age, years ± SD | 46 ± 15 | 42 ± 13 | 0.23 | 44 ± 14 |
| Female sex, N (%) | 18 (47) | 13 (50) | 0.84 | 33 (52) |
| Race/Ethnicity, N (%) |  |  | 0.06 |  |
| Caucasian | 32 (84) | 19 (73) |  | 51 (80) |
| Hispanic | 2 (5) | 0 (0) |  | 2 (3) |
| African American | 0 (0) | 3 (12) |  | 3 (5) |
| Asian | 0 (0) | 2 (8) |  | 2 (3) |
| Other | 4 (11) | 2 (8) |  | 6 (9) |
| Average BMI ± SD,<br>kg/m<sup>2</sup> | 26.9 ± 7.5 | 25.8 ± 5.4 | 0.49 | 26.5 ± 6.7 |
| Disease phenotype, N<br>(%) |  |  |  |  |
| Fistulizing | 9 (24) | 8 (31) | 0.53 | 17 (27) |
| Fibrostenotic | 7 (18) | 7 (27) | 0.42 | 14 (22) |
| Disease Location, N<br>(%) |  |  | 0.41 |  |
| Ileum only | 12 (32) | 5 (19) |  | 17 (27) |
| Colon only | 5 (13) | 6 (23) |  | 11 (17) |
| Ileum and colon | 21 (55) | 15 (58) |  | 36 (56) |
| Concomitant<br>Medication Use, N (%) |  |  |  |  |
| Immunomodulator | 28 (74) | 20 (77) | 0.77 | 48 (75) |
| Corticosteroids | 28 (74) | 18 (69) | 0.70 | 46 (72) |
| Prior anti-TNF<br>exposure | 32 (84) | 24 (92) | 0.34 | 56 (88) |

The probability curve of clinical response to ileal MVL was bell-shaped, with a range of 1.35-1.55 µm associated with a significantly higher response rate of 82% (14/17), compared to 51% (24/47) for values outside this range (p=0.04, **Figure 3A**). Pre-treatment ileal MVL did not correlate with either baseline clinical disease activity by HBI (rho = −0.12) (**Figure 3B)**, or baseline endoscopic disease activity by SES-CD (rho = - 0.28) (**Figure 3C**).

**Figure 3.**
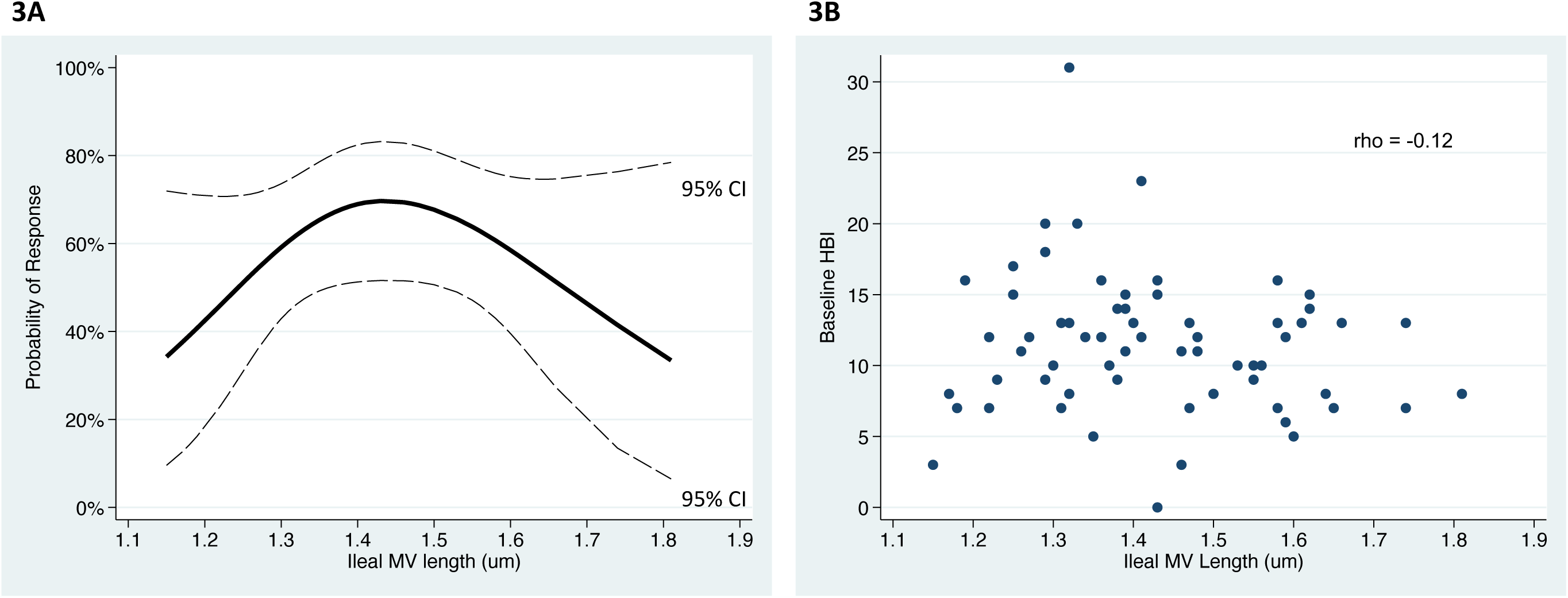
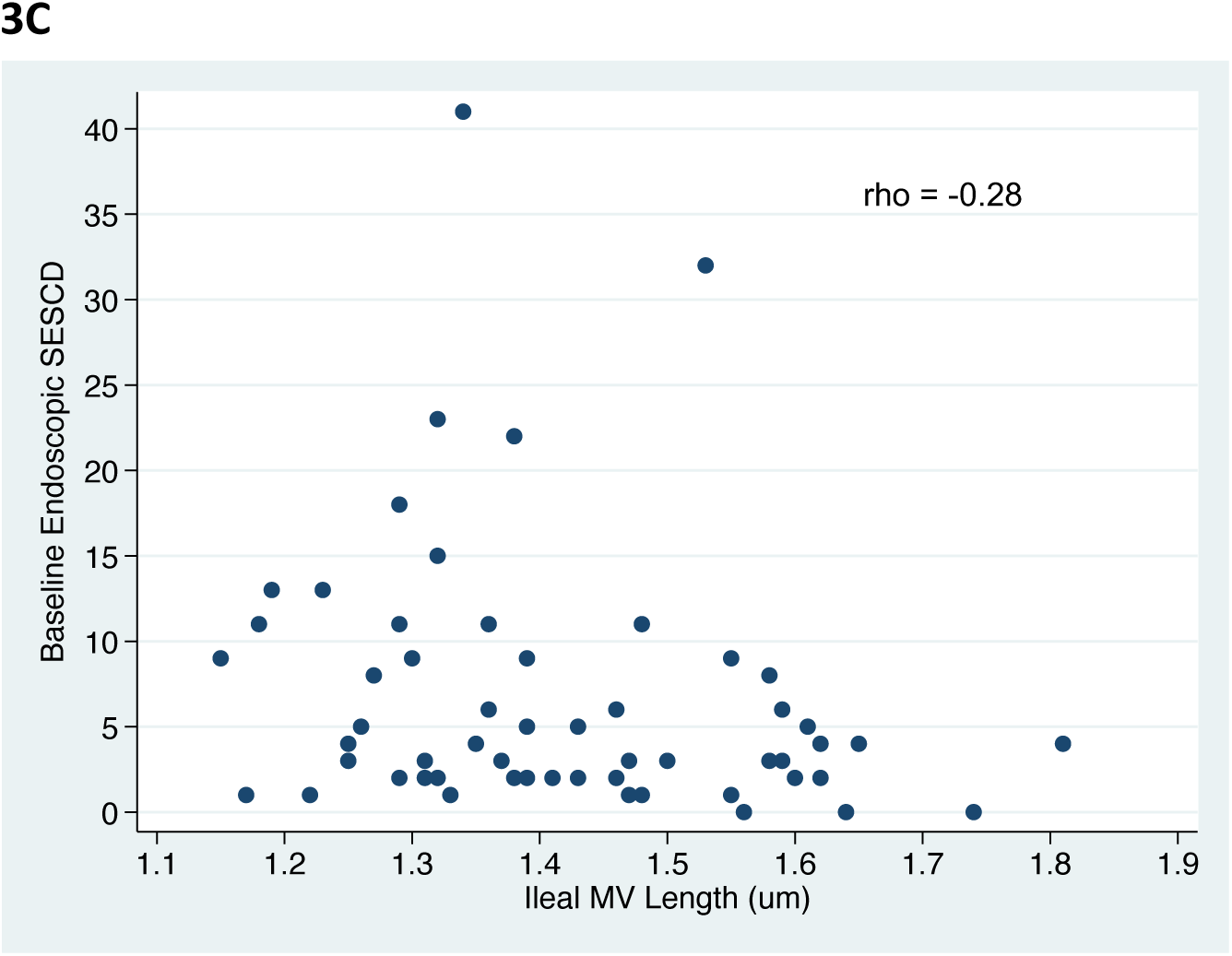
Ileal MVL is predictive of response to vedolizumab in CD patients from the multicenter retrospective cohort. A) Clinical response as a function of ileal MVL (N=64). B) Correlation between ileal MVL and baseline clinical disease activity by HBI (N=64). C) Correlation between ileal MV length and baseline endoscopic disease activity by SES-CD (N=58).

Pre-treatment ileal MVL also did not correlate with pre-treatment ileal IEC pyroptosis (rho = 0.01) (**Figure 4A)**. Moreover, the combination criteria of ileal IEC pyroptosis level < 14 positive cells / 1,000 IECs and ileal MVL of 1.35-1.55 µm was associated with a vedolizumab clinical response rate of 81% (25/31), compared to 39% (13/33) for those with neither criterion (p < 0.001); corresponding rates of clinical remission rates were 55% (17/31) vs. 24% (8/33) in the 2 groups (p=0.012), respectively (**Figure 4B)**. The combination criteria could identify 66% of responders (25/38), compared to 42% of responders using ileal IEC pyroptosis alone and 37% of responders using MVL alone. Additionally, fulfillment of neither criterion identified the majority (77%, 20/26) of non-responders to vedolizumab.

**Figure 4.**
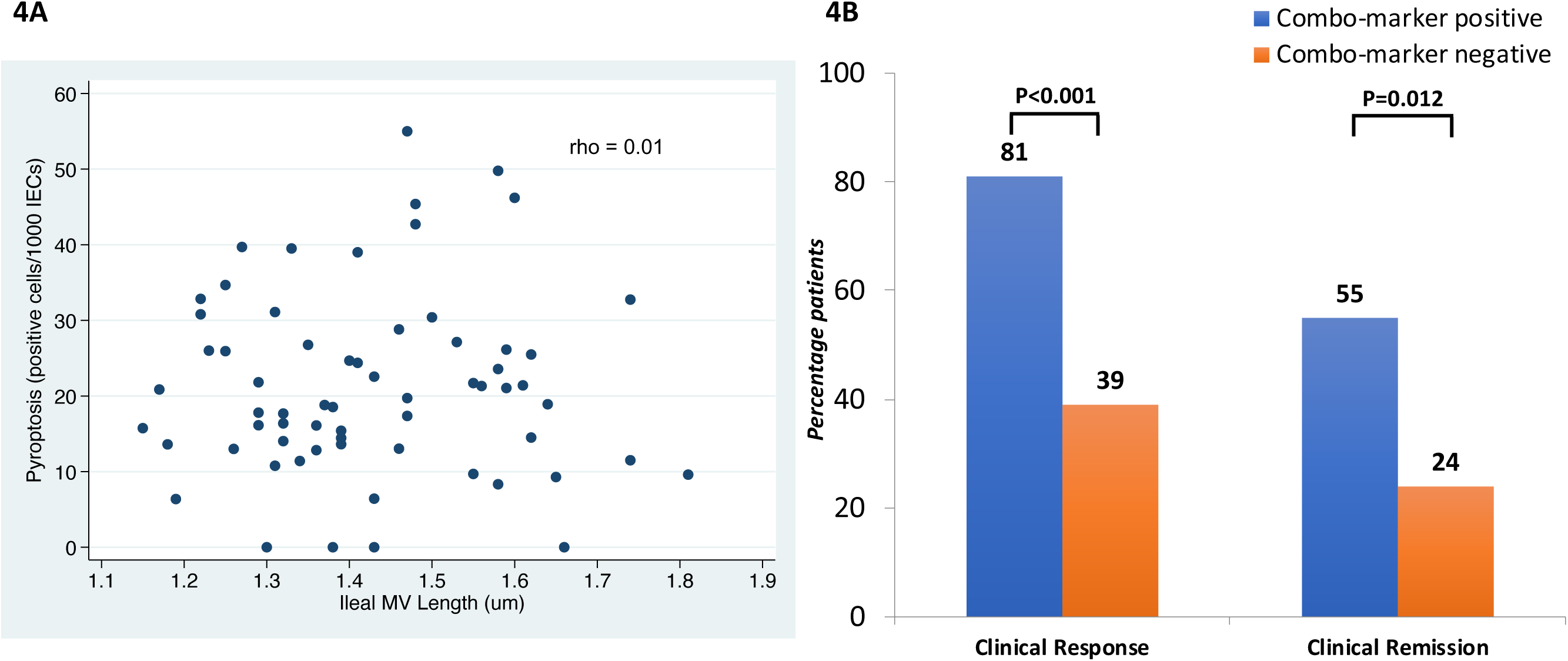
Ileal MVL and ileal IEC pyroptosis are independent and synergistic biomarkers for prediction of response to vedolizumab in CD. A) Correlation between ileal MVL and ileal IEC pyroptosis on pre-treatment biopsy samples (N=64). B) Improved discriminant ability for responders from non-responders to vedolizumab (N = 64).

## Discussion

At the present time, selection of biologic therapy in IBD is largely arbitrary. The transformational goal of future IBD therapy is the personalized selection of agents based on each individual patient’s pathobiology. In this study using both prospective randomized and multicenter retrospective cohorts of CD patients, ileal MVL, an epithelial biomarker, was found to be predictive of response to two different biologic agents, ustekinumab and vedolizumab. Additionally, combining this mucosal biomarker with another recently discovered biomarker, ileal IEC pyroptosis, was associated with high rates of clinical response to vedolizumab in CD and also identified the majority of non-responders to vedolizumab. These findings are consistent with our hypothesis that epithelial cellular function may play an important role not only in the pathogenesis of IBD but also in guiding the selection of biologic agents for individual CD patients.

Interestingly, the probability curves for clinical response by ileal MVL for the 2 biologic drugs evaluated were different: a linear relationship for ustekinumab and a bell-shaped curve for vedolizumab. The biological mechanism by which ileal MVL is associated with response to both ustekinumab and vedolizumab has yet to be elucidated but is being actively explored by our group. As was observed with ileal IEC pyroptosis in vedolizumab-treated CD patients,^21^ pre-treatment ileal MVL did not correlate with either baseline clinical or endoscopic disease activity in both ustekinumab-treated and vedolizumab-treated CD patients. Additionally, these 2 mucosal biomarkers at pre-treatment did not correlate with each other and seem largely independent in terms of utility to predict response to vedolizumab in CD patients with effects that were nearly additive. Also under current investigation by our group is whether ileal IEC pyroptosis is associated with response to ustekinumab in CD.

There is an urgent and unmet need in the care of patients with moderate-to-severe CD to personalize biologic therapy. The existing CD phenotypic classification system which focuses on inflammation and complications is not sufficient to guide treatment decisions for biologics, which are designed to target specific inflammatory pathways. Due to limited therapeutic efficacy and lack of personalized treatment, IBD-related healthcare outcomes and costs have not improved with the widespread clinical adoption of biologic therapies over the past 2 decades. The annual costs of IBD treatment have increased by 30% within the last 5 years, and IBD-related hospitalizations quadrupled from 1998 to 2015.^10, 11^ Nearly half of CD patients placed on a given biologic therapy do not have a clinical response, likely because the therapy chosen is not targeting the specific inflammatory pathway(s) driving the disease.

To address this need, some recent effort has focused on the discovery of biomarkers that can predict therapeutic responses to various biologics. Several mucosal biomarkers have been reported to be associated with response to anti-TNF agents in CD: oncostatin M,^15^ TNFR2 and IL13RA2, ^16, 17^ and TREM1.^18, 19^ For example, high pre-treatment levels of transmembrane TNF positive cells were observed to be positively associated with response to adalimumab,^22^ whereas high pre-treatment levels of oncostatin M were found to be negatively associated with response to infliximab and goliumumab.^19^ With respect to predicting response to anti-integrin therapy, high mucosal gene expressions of αE^23^ and granzyme A^24^ were demonstrated to be associated with improved response to etrolizumab in patients with UC.

Over the past decade, advances in disease pathogenesis from multi-omics approaches have identified that epithelial barrier function is responsible for ongoing mucosal inflammation in IBD.^12-14^ Advances in innate immunology by our group have identified ileal IEC pyroptosis as a possible contributor to mucosal barrier dysfunction and inflammation in IBD.^25^ Our group recently reported that lower levels of ileal IEC pyroptosis were associated with increased clinical response and remission to vedolizumab in patients with CD.^21^ Based on these emerging data, we investigated another mucosal biomarker that may play a key role in epithelial cell biology and function, ileal MVL, as was recognized to be a measure of cellular absorption and metabolism.^26^ The results of the current study extend the use of MVL to enhance our ability to predict response to vedolizumab in CD, with additive and synergistic effects of the combination of the two independent biomarkers. Additionally, this study was able to extend biomarker-assisted prediction of clinical and endoscopic outcomes to ustekinumab-treated patients.

The strengths of this study also include the use of prospective data from a pivotal randomized controlled trial of ustekinumab in CD. Additionally, for the retrospective cohort study, data were pooled from 5 high-volume IBD centers, thus conferring some validity to the data collection and a degree of variation to the patient population. This study also had several limitations. Follow-up endoscopic data in the randomized cohort were not available for all patients, and many of the patients in the retrospective cohort did not have follow-up colonoscopies at the time of assessment of clinical response and remission. Also, in the retrospective cohort, many patients did not have complete data on pre- and post-treatment biochemical markers, such as CRP and fecal calprotectin, and thus these were not studied. Finally, our findings may not be generalizable to patients who are ineligible for clinical trials (for the ustekinumab inquiry) or those from more community settings (for the vedolizumab inquiry).

In conclusion, we have found that a previously identified mucosal biomarker, ileal MVL, could predict response to both ustekinumab and vedolizumab in CD. Combining this biomarker with another recently identified mucosal biomarker, ileal IEC pyroptosis, enhanced the association with response to vedolizumab in CD. Future larger studies validating our data are warranted. Additionally, future studies investigating the clinical significance of both biomarkers for prediction of response to vedolizumab and ustekinumab in UC as well as for prediction of response to anti-TNF agents in both CD and UC are planned from our group.

## Acknowledgement

We would like to thank Takeda Pharmaceutical for their funding support of this study.

